# Stepwise Isolation of Diverse Metabolic Cell Populations Using Sorting by Interfacial Tension (SIFT)

**DOI:** 10.1101/2024.09.23.612740

**Authors:** Matthew Shulman, Thomas Mathew, Aria Trivedi, Azam Gholizadeh, Charlotte Colcord, Ryan Wiley, Kiron S. Allen, Lakshmi Thangam, Kelsey Voss, Paul Abbyad

## Abstract

We present here a passive and label-free droplet microfluidic platform to sort cells stepwise by lactate and proton secretion from glycolysis. A technology developed in our lab, Sorting by Interfacial Tension (SIFT), sorts droplets containing single cells into two populations based on pH by using interfacial tension. Cellular glycolysis lowers the pH of droplets through proton secretion, enabling passive selection based on interfacial tension and hence single-cell glycolysis. The SIFT technique is expanded here by exploiting the dynamic droplet acidification from surfactant adsorption that leads to a concurrent increase in interfacial tension. This allows multiple microfabricated rails at different downstream positions to isolate cells with distinct glycolytic levels. The device is used to correlate sorted cells with three levels of glycolysis with a conventional surface marker for T-cell activation. As glycolysis is associated with both disease and cell state, this technology facilitates the sorting and analysis of crucial cell subpopulations for applications in oncology, immunology and immunotherapy.

## Introduction

Modern cell biology and medicine rely on robust methods to separate distinct cell populations for downstream usage or analysis. The workhorse of cell sorting is Fluorescence-activated cell sorter (FACS), where cells are sorted based on signal from a specific marker, typically a fluorescent probe. Many applications require separations beyond binary populations and modern FACS instruments can sort up to six populations. Although expensive (multiplexed FACS instruments are upwards of 200K), the specificity of FACS is unmatched by other techniques. However, FACS analysis is fundamentally limited by both the availability and selectivity of probe molecules. It also is not amenable to sorting directly based on cell secretions or metabolic activity.

Label-free microfluidic techniques, such as inertial microfluidics,^1,2^ acoustophoresis,^3^ dielectrophoresis^4^ and deterministic lateral displacement ^5,6^ provide ease of use and high throughput in sorting. They are particularly well suited to sorting into three or more populations as this capacity can be integrated with little added expense and complexity by adding channel outlets at different lateral positions.^3,7,8^

Most cellular processes, especially metabolism, function on a continuum rather than in binary on-off states. Instead, they are finely tuned to varying levels of activity to meet the needs of the cell. Thus, quantitative measurements of metabolism are routinely used for both fundamental research and the study of disease states in particular cancer and metabolic disorders. To emphasize this, extracellular flux analyses (EFA), which measures metabolism through oxygen consumption and extracellular acidification in bulk cells, is referenced in over 7,000 publications. A hallmark of cancer cells is their aberrant metabolism^9^ which is used for both cancer diagnostics and monitoring. Glycolysis, measured by extracellular acidification, can serve as a biomarker to isolate tumor cells that do not have general markers.^10^ This metabolic reprogramming to favor glycolysis over oxidative phosphorylation is a shared feature of many fast dividing cells such as cancer cells, activated T-cells and stem cells.^11,12^ Moreover, cell metabolism is now considered a main driver and decision point for cell proliferation, differentiation and disease progression.^13^ Target cell populations do not always have the highest metabolism. As one example, the level of glycolysis in activated T-cells regulates their differentiation, with intermediate glycolytic activity^14^ (between that of naive and helper T-cells) steering differentiation toward rare, therapeutically valuable regulatory T- cells (Treg). In another setting, long-lived memory T-cells can be distinguished from effector T-cells based on lower levels of glycolysis.^15,16^ This highlights the need for techniques to isolate cells with a progression of glycolytic profiles.

Our lab has developed the droplet microfluidic technique, <u>S</u>orting by Inter<u>f</u>acial <u>T</u>ension (SIFT), that allows for the passive and label-free sorting of droplets by pH. It has been used to sort enzymes,^17^ cells^18–20^ and amplified DNA.^21^ The technique exploits a dependence of droplet interfacial tension on pH. Cellular glycolysis involves the secretion of protons, which lowers the pH of droplets. This enables the passive selection of droplets by interfacial tension and hence single-cell glycolysis. The key feature of the work presented here is that the droplets undergo two successive and distinct changes in pH in the sorting device. The first is due to the secretion of protons from cell glycolysis.

The second is due to the adsorption of surfactant onto the droplet interface. As this second change in pH is dynamic, the interfacial tension also progressively increases as the droplets flow downstream. In this paper, we first characterize the change in pH due to the adsorption of surfactant. To characterize the full range of droplet pH, we integrate three different pH determinations, one colorimetric and two based on fluorescence. We then utilize multiple rails positioned at various points along the channel to select cells with decreasing glycolysis levels. Cells are guided to different chip outlets and collected using a protocol designed for recovery and viability.

The separation of activated T-cells with different metabolic levels is particularly important, as glycolysis is not merely a downstream effect but actually a key regulator of cellular differentiation outcomes.^14,22^ We have previously shown that SIFT can be used to separate highly activated cells from naive cells.^23^ Here, for the first time, the single-cell measurements of three levels of glycolysis are correlated with conventional activation markers for T-cells. This assay demonstrates the device’s potential to make new cell correlations and isolate multiple cell subpopulations with distinct glycolysis profiles.

## Materials and Methods

### Cells

K562 human chronic myelogenous leukemia cells were grown in ATCC-formulated Iscove’s Modified Dulbecco’s Medium (IMDM) and Jurkat Clone E6-1 TIB-152™ Human Acute T-cell leukemia cells were grown in ATCC-formulated RPMI-1640 Medium. Both cell lines were purchased from ATCC. Cells were grown at 37 °C in a 5% CO_2_ atmosphere supplemented with 10% fetal bovine serum (HyClone, GE Healthcare Life Sciences, Logan, UT) and 2% v/v penicillin-streptomycin (10 000 units/mL-10 000 μg/ mL) solution (Gibco, Life Technologies Corporation, Grand Island, NY).

### Cell Activation and Preparation for on-chip experiments

Jurkat cells were activated with soluble activation complexes (ImmunoCult, StemCell Technologies) following manufacturer protocol and incubated for 24 hours. On the day of the experiment, activated Jurkat T-cells were centrifuged, washed and resuspended in HBBS. To better identify droplet occupancy, cells were labeled by incubating for 30 min at 37 °C and 4% CO_2_ atmosphere with Calcein AM (Thermo Fischer, Waltham, MA, USA), a viability fluorescent dye. Calcein was not used when staining with CD69 to avoid spectral overlap in the fluorescence signals. Subsequently, the cells were washed again and resuspended in on-chip solutions at a cell concentration of 5 X 10^5^ cells/mL, which was determined using a Cellometer Auto T4 Bright Field Cell Counter (Nexcelcom Bioscience LLC, Lawrence, MA) to ensure single cell occupation of droplets. On-chip solutions were a 1:1 mix of media and 1.5 mM PBS buffer. The media was prepared without fetal bovine serum (deproteinated media), both solutions were supplemented with 1% w/w Pluronic F-68 (Affymetrix Inc., Maumee, OH), 15% v/v Optiprep solution (Fresenius Kabi Norge AS for Axis-Shield PoCAS, Oslo, Norway) and 0.1 mg/mL pyranine (AAT Bioquest Inc., Sunnyvale, CA). Solution pH and osmolality (determined with Vapro Vapor Pressure Osmometer 5520, Wescor, ELITech Biomedical Systems, Logan, UT) of on-chip solutions were adjusted to physiological values (pH 7.4-7.6; 280−320 mOsmol) prior to experiment. Pluronic F-68 was used to promote droplet stability and cell viability, whereas Optiprep modulated solution density to limit cell sedimentation within the tubing and droplets. Pyranine (0.1mg/mL, AAT Bioquest Inc., Sunnyvale, CA) or fluorescein disodium salt (0.02mg /mL, Eastman Kodak Co., Rochester, NY) were added to the cell solutions in some experiments to serve as a fluorescent ratiometric pH probe for determining droplet pH on chip.

### CD69 Antibody Assay

The cells were prepared as described above. After activation, the cells were incubated in media at 37 °C in a 5% CO_2_ atmosphere for 24 hours to allow for the display of CD69 surface markers specific to activation. Cells were washed and resuspened in HBSS before staining with Human CD69 APC conjugate (Thermo Fisher Scientific). Staining was performed at a v/v ratio of cell suspension (∼1X10^6^ cells/mL) to CD69 staining solution of 100:1 for 30 minutes at room temperature. The cells were then centrifuged, washed with HBSS, and resuspended in on-chip solutions before being injected into the microfluidic chip.

### Microfluidic Device

Chips with channel depth modulations were fabricated from polydimethylsiloxane (PDMS), utilizing the dry-film photoresist soft lithography technique previously reported by Stephan et al.^24^ This technique facilitates easy prototyping with multilevel designs. The PDMS chip was irreversibly bonded to a glass slide via plasma treatment. To render the internal surfaces of the channel hydrophobic, the channels were treated with Novec 1720 electronic grade Coating (3M, Maplewood, MN) for 30 min at 150 °C. The channel design, rail position and rail dimensions of the device to sort three populations of cells are provided in Supporting Information (SI) Figures S1, S2, and S3 respectively.

### Droplet Sorting and Measurements

The general use of the sorting device was similar to what has been described previously.^18,23^ Briefly, the chip consists of a droplet generator where cells are encapsulated into droplets; an incubation channel enabling a change in droplet pH due to the cells’ metabolism; and a sorting region. Cellular solution was injected into the chip through an aqueous inlet. Via a flow focuser, droplets were generated in 0.1% w/w Picosurf-1 surfactant oil (Sphere Fluidics Limited, Cambridge, United Kingdom) in Novec 7500. An additional oil outlet after droplet generation was set to flow in the opposite direction of the main flow to reduce the amount of oil before the droplets entered the incubator region. This enabled tight packing of droplets within the incubator to ensure the same incubation times for all droplets.^25^ The length of the incubation channel was 20 cm, double the length used previously.^18,23^ Average incubation time ranged from 10 to 20 minutes depending on the experiment before the channel narrowed and droplets entered the sorting region. At the end of the incubator, oil solution, QX100 droplet generation oil for probes (Biorad, Hercules, CA), entered the chip through two inlets, the QX100 inlet and the Oil Entrainment Inlet (Figure S1). This oil/surfactant combination is called here QX100 for simplicity and consistency with prior publications. Droplets entered the sorting region that included one or more rails, of higher channel height (Figure S2). The rails, oriented at 45 degrees to the flow direction, allowed sorting droplets by interfacial tension and hence pH.

Flows within the chip were controlled via a computer-controlled syringe pump system (Nemesys, Cetone, Korbussen, Germany). Typical flow conditions can be found in the Supplemental Table S1. The temperature of the chip during experiments was maintained at 37 °C using a heating stage with a control module and temperature feedback (CHS-1 heating plate, TC-324C temperature controller, Warner Instruments, Hamden, CT).

On-chip images and videos were taken on an inverted fluorescence microscope (Olympus IX-51) equipped with a 4X objective, a shuttered LED fluorescence excitation source (Spectra-X light engine, Lumencor, Beaverton, OR) and a high-speed camera (VEO-410, Vision Research, Wayne, NJ). The microscope filter cube contained a dual-edge dichroic mirror (Di03-R488/561-t1-25 × 36, Semrock, IDEX Health & Science LLC Rochester, NY) and dual-band emission filter (FF01−523/610-25, Semrock) that enabled transmission of pyranine, fluorescein and Calcein AM fluorescence. An excitation source with individually addressable LEDs coupled to an Arduino (Arduino LLC, Scarmagno, Italy) for rapid alternation between different colored LEDs using simple TTL triggering was used to determine droplet pH values. Droplets were excited with alternating violet (395 nm BP 25 nm), blue (440 nm BP 20 nm) and green excitation (561 nm BP 14 nm) at a rate of 100 frames per second (33 fps for each color) for pyranine pH measurements. For long sorting experiments, 2 minute videos were taken about every 10 minutes and data was combined. For fluorescein pH measurements, droplets were excited with alternating blue, violet and cyan (479 nm BP 34 nm) excitation light. Color images were obtained on an inverted fluorescence microscope (Olympus IX-50) equipped with a digital SLR camera (Canon EOS 70D).

### Cell Collection

The general workflow for cell collection is summarized in Figure S4. Cells were first sorted as described above and collected into 1mL pipette tips inserted directly into the chip outlets. The pipette tips were prefilled with 0.5 mL of Novec 7500 to dilute the surfactant found in QX100. Minimizing exposure to this surfactant was found to improve cell viability. 300 ul of HBSS droplets (diameters of 50-200 μm) made in 0.1% w/w Picosurf-1 surfactant oil was also added to the pipette tips. The empty droplets improved cell recovery by ensuring that sorted cell droplets didn’t collect on the pipette walls and by facilitating droplet coalescence. To avoid overflowing the pipette tip over the course of the experiment, oil was intermediately removed from the pipette tip using a long blunt needle syringe. Care was taken during the removal of oil to avoid disturbing the chip or provoking the coalescence of the droplets layered above the oil. After each oil removal, 0.5 mL of Novec 7500 was added to the pipette tip.

At the end of the sorting experiment, pipette tips were removed and the oil was drained from the bottom of the tip. Droplets were then collected into microcentrifuge tubes. Droplets were coalesced using a static gun (MILTY Pro Zerostat 3)^26^ and transferred into a 96 well fluorescence plate. To sediment the cells to the bottom of the plate for imaging, the plate was centrifuged at 1500 rpm (g-force 525) for 5 minutes using a TX-100 swinging bucket rotor (Thermo Scientific) in a Sorvall Legend XTR Centrifuge.

### Data Analysis

ImageJ software was used for image analysis.^27^ pH values of individual droplets were determined at the end of the incubation channel before droplets entered the sorting region via the ratio of fluorescence intensity from background-subtracted ratio of two color excitation. For pyranine, a calibration curve from fluorescence ratios of blue and violet excitation for droplets of known pH was used to determine pH using a procedure described previously.^19^ For fluorescein, a similar procedure was used based on the ratio of fluorescence of cyan and blue excitation (calibration curve provided in Figure S5). Green excitation was used to identify cells labeled with Calcein AM. Logistic regression was used to statistically estimate optimal pH thresholds to separate selected from non-selected cells. The pH threshold was defined at a 50% predicted probability of selecting the cell. The standard error of the prediction was used to obtain a 95% confidence interval around that threshold.

Collected cells were imaged with an inverted fluorescence microscope (Olympus IX-50) equipped with a 20X objective, a shuttered LED fluorescence excitation source (Sola SE-II), Lumencor, Beaverton, OR) and a CMOS camera (Orca Flash 2.8, Hamamatsu). Fluorescence images were obtained with long exposure (1.9 seconds) using a red fluorescence cube (Ex: 631 nm BP 28 nm, Dichroic mirror 652 nm, Em: 680 nm BP 42 nm). Debris and irregularly shaped cells were excluded from the analysis. Average cellular fluorescence was measured in ImageJ.

## Results and Discussion

SIFT allows for the passive and label-free sorting of cells based on cell glycolysis. The technique leverages a dependence on the droplet interfacial tension with pH in the presence of a surfactant. Cell glycolysis is coupled with a secretion of protons, leading to a decrease in droplet pH. This enables the selection of droplets by interfacial tension and hence single-cell glycolysis.

Figure 1 presents the chip geometry and technique (Figure 1a) with a pH indicator to provide a direct visualization by color of the droplet pH at the three sequential steps on the device: cell encapsulation, incubation and sorting. This figure illustrates a single rail; however, later sections of this study will include devices featuring multiple rails. A supplemental video pans the entire chip in the flow direction (Video S1). Images and video were captured with a commercial SLR color camera. The pH color indicator, phenol red, was added to the cell media and undergoes a color change from red to yellow from pH 7.5 to 6.9 (color scale provided in top right). Phenol red is biocompatible and is commonly used in growth media to monitor pH.

**Figure 1.**
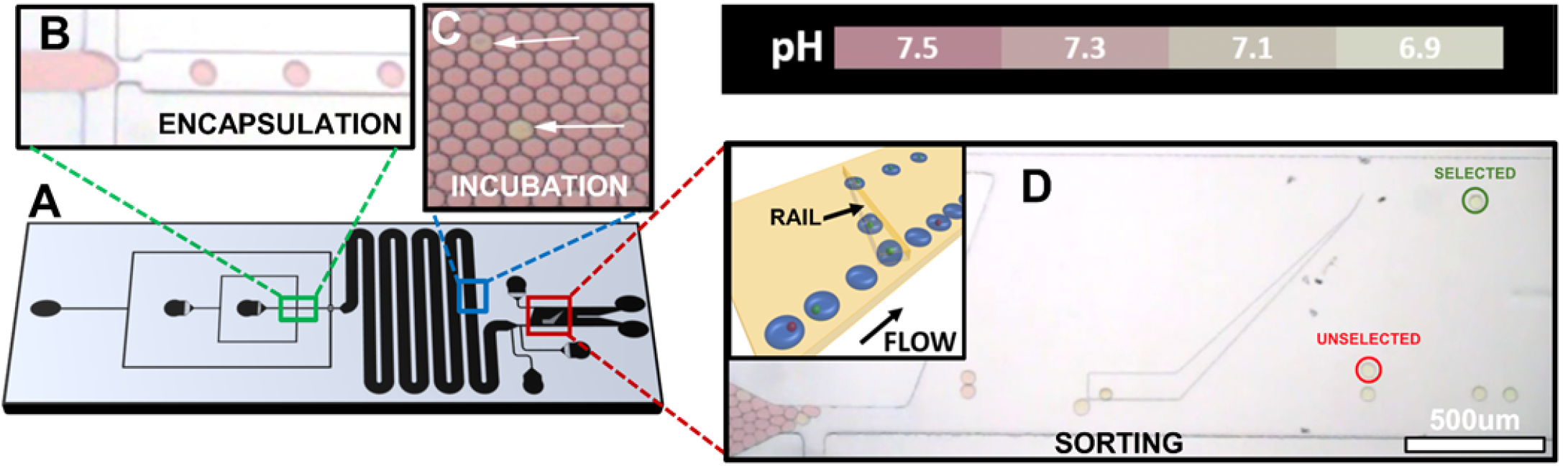
Design and color images of SIFT a) Channel geometry b) Droplet formation and cell encapsulation c) Droplets after 10 min. of incubation. Arrows point to orange droplets of lower pH due to cell metabolism. d) Droplets containing cells with high metabolism ride the rail laterally up (circled in green). Empty droplets or those containing cells with low metabolism (circled in red) are only slightly deflected by the rail. Inset shows droplet deformation in rail. pH color scale shown in top right.

Cells are encapsulated into droplets (Figure 1b). Cell density is kept low to avoid double occupancy of cells in droplets. Typically only one in twenty or thirty droplets contain a cell. Upon encapsulation, the pH of all droplets is unchanged from the media preparation and is around a pH of 7.5. Droplets flow through a long serpentine incubation channel. The majority of droplets are empty and will remain unchanged throughout the length of the incubator (Figure 1c). However, encapsulated cells will secrete protons via glycolysis and reach a lower droplet pH indicated by a color change from red to shades of orange (arrows Figure 1c). Cells have heterogeneous glycolysis activity and the number of droplets that show a clear distinction in color increases further along in the incubation channel.

The length of the incubator channel was doubled to 20 cm compared to the channel designs reported previously.^18,23^ To obtain a significant color contrast for the thin pathlength (25 μm channel height), phenol red was dissolved at the relatively high concentration of 2.2 mM. At this concentration, the indicator itself is an important contributor to the buffer of the media solution that would be 1.5 mM in the absence of the phenol red. The longer incubation channel allows incubation times of 10-20 minutes which was sufficient to ensure a noticeable color change in droplet pH before reaching the sorting region. This longer incubation channel was used in all experiments presented in this paper.

After incubation, droplets enter the sorting region where carrier oil containing an acidic surfactant is introduced into the chip. In the presence of this surfactant, droplets exhibit an inverse dependence of pH and interfacial tension. This oil/surfactant combination, Droplet Generation Oil for Probes (Bio-Rad), is called here QX100 for simplicity and consistency with prior publications. In the sorting region droplets encounter a microfabricated trench, or rail. Droplets are pancake shaped, confined by the top and bottom of the channel. The droplets become less confined in the rail, lowering their overall surface area (Figure 1d, inset).^28^ The red droplets at high pH (low interfacial tension), enter the rail, but are pushed off by the entrainment flow of the oil, directed towards the Unselected outlet (Figure 1d, circled in red). These red droplets would contain no cells or cells with low glycolysis. In contrast, when an orange droplet at low pH, hence high interfacial tension, enters the rail, the entrainment flow is insufficient to push the droplets from the rail. These orange droplets, containing cells with high glycolysis, are displaced laterally by the rail and are directed towards the Selected outlet (Figure 1d, circled in green). Hence cells are sorted based on the biological acidification from glycolysis that occurs in the incubation channel. The method has good selectivity, separating droplets with differences of 0.05 pH units.^19^ However, to date the method was limited to binary sorting.

The key to the presented work is that all droplets in the sorting region exhibit a *<u>second</u> <u>acidification</u>* due to the adsorption of the new acidic surfactant.^29^ This can most easily be discerned from the progressive color change to yellow of all droplets as they flow in the sorting region (Figure 1d). The pH of a droplet when it encounters the rail is thus an additive effect of both effects, acidification from glycolysis and surfactant adsorption.

The phenol red indicator allows direct observation of the acidification, however it is difficult to make quantitative pH readings based on color. Moreover, the color change only captures part of the pH change. Phenol red has a pKa of 7.7.^30^ Below 7.0, all droplets are yellow in color and cannot be distinguished. We have used pyranine as a ratiometric fluorescence pH sensor to determine droplet pH. With a pKa of around 7.3,^31^ a pH between 6.8 and 8.0 can be measured. This range is ideally suited to measure the biological acidification from glycolysis. However, it only captures part of the acidification from surfactant adsorption that results in a droplet pH below 6.5 (Figure S6). Fluorescein’s emission intensity is also modulated by pH and has been previously used as a ratiometric pH sensor.^32^ With a pKa of 6.4,^33^ around a unit below pyranine, it can measure pH between 5.8 and 7.6. It is thus well-suited to measure the acidification from surfactant adsorption and is used to determine the kinetics of the acidification in the sorting region.

The second acidification is time dependent with droplets attaining lower pH as they flow downstream. Figure 2a and 2b show fluorescence images with cyan excitation of a series of droplets containing fluorescein as they flow in the sorting region. These measurements were performed on chips with no rail in the sorting region to avoid obstructing the images. The far left of the image is the end of the incubation channel and tapered. QX100 enters the device by a channel on the bottom left of the image. The fluorescence of fluorescein diminishes with a decrease in pH. The droplet’s fluorescence intensity decreases as they flow to the right, thus consistent with a decrease in droplet pH from adsorption of the acidic surfactant. Moreover, this pH change is not observed when the QX100 is replaced with a non-ionic picosurf surfactant in the sorting region. Figure 2a and 2b show images of droplets with initial droplet pH values of 7.56 and 6.98, respectively. Although both show a progressive decrease in fluorescence as droplets flow right, the droplets with an initial pH of 6.98 exhibit both lower initial fluorescence and a more pronounced reduction in brightness at downstream positions.

**Figure 2.**
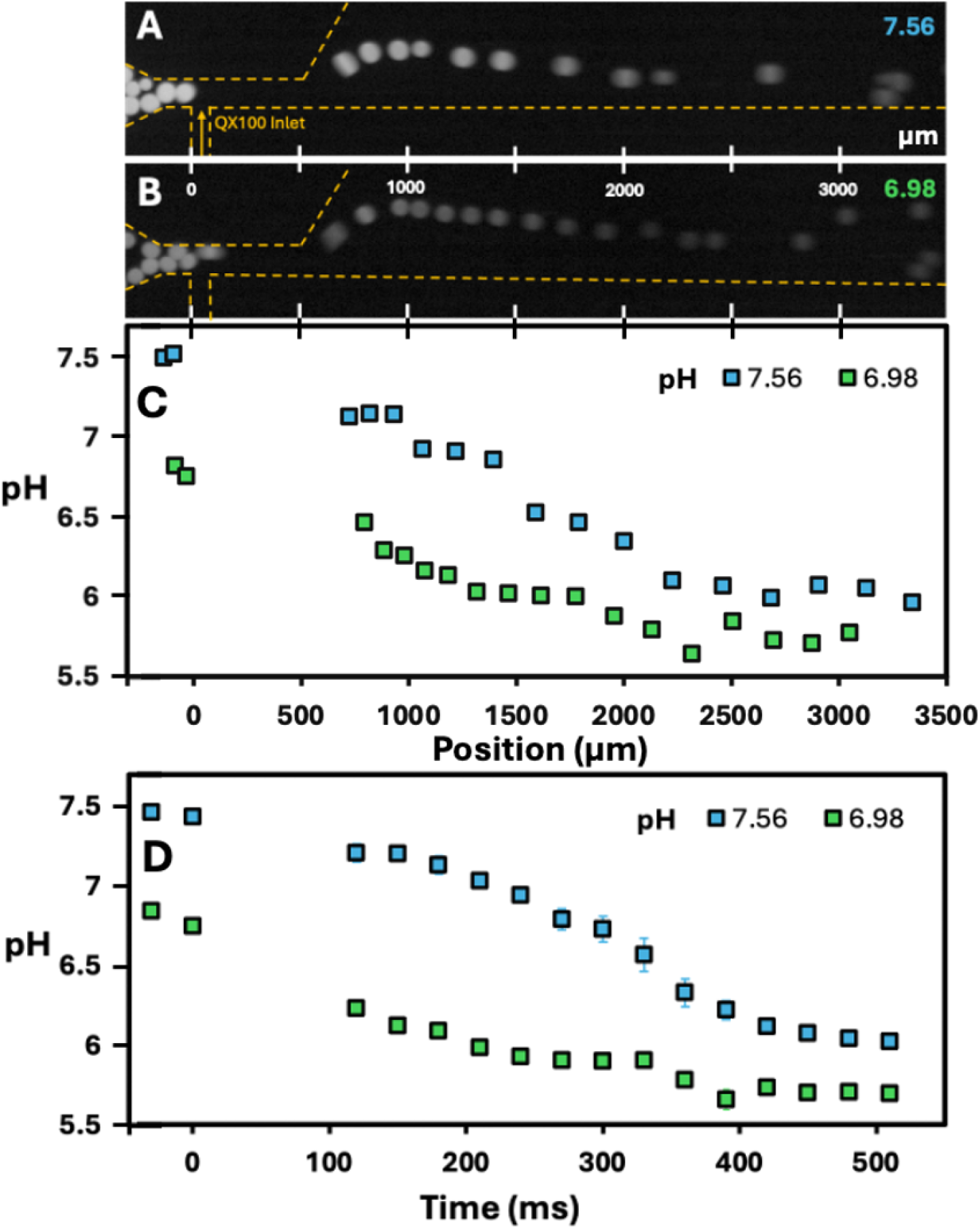
Acidification of droplets in the sorting region from adsorption of an acidic surfactant. a) Droplets with a higher initial pH (7.56) exposed to surfactant visualized under cyan excitation. A greater intensity in the cyan channel indicates a higher pH. b) Droplets with a lower initial pH (6.98) exposed to surfactant. c) Representative pH of a single flowing droplet as a function of position as determined by fluorescent measurements of fluorescein, a ratiometric pH probe. d) Average (± SEM) pH of flowing droplets as a function of time after exposure to surfactant (n=5).

Using a calibration curve (Figure S5), droplet pH can be determined as a function of position and time (Figure 2c and 2d). The zero position and time here are defined by the entry of the QX100 channel, with negative values upstream of the QX100 channel. Figure 2c shows the pH of a representative droplet as it flows from left to right with the position scale directly matching the images in Figure 2a and 2b above. pH readings were not possible for the first 600 μm due to the fast movement of the droplets as they flow through the narrow channel. The pH value of droplets depends on the droplet pH when entering the sorting region. For example, for the initial pH of 6.98 the droplets will attain a pH of 6.25 at around 1000 μm. The droplet at an initial pH of 7.56 will only attain this same pH at around 2000 μm. The droplet pH can also be followed as a function of time (Figure 2d), with a change in pH occurring within the first hundreds of milliseconds after contact with the QX100. The acidification appears faster than reported in a previous study,^34^ where the adsorption of carboxylic acid surfactant to a droplet interface occurred over a few seconds. However, a direct comparison is complicated because of the differences in surfactants and the fact that the previous study examined adsorption at a bare interface. The spatial and temporal pH data display a similar profile. In comparing the different initial pH, the low initial pH of 6.98 displays a lower pH at all time points. As pH has been correlated to interfacial tension with QX100,^18^ this dynamic acidification leads to a concurrent rise in droplet interfacial tension. Importantly, this increasing interfacial tension can be leveraged using rails to select droplets at different downstream positions.

To characterize droplet sorting at different downstream positions, a microfluidic chip was designed with six rails, each separated by 300 μm (Figure 3a). K562 cells were encapsulated into droplets and incubated for approximately 14 minutes. These cells were used as their high glycolysis ensured a large spread of droplet pH values. At the end of the incubation channel, empty droplets remained near the initial pH of 7.4, while droplets containing cells ranged from a pH of 6.8 to 7.4. Droplets were selected by different rails (Video S2), numbered from left to right based on downstream position. Figure 3a shows the selection of droplets by rail 1, rail 6 and an unselected droplet circled in green, blue and red respectively.

**Figure 3.**
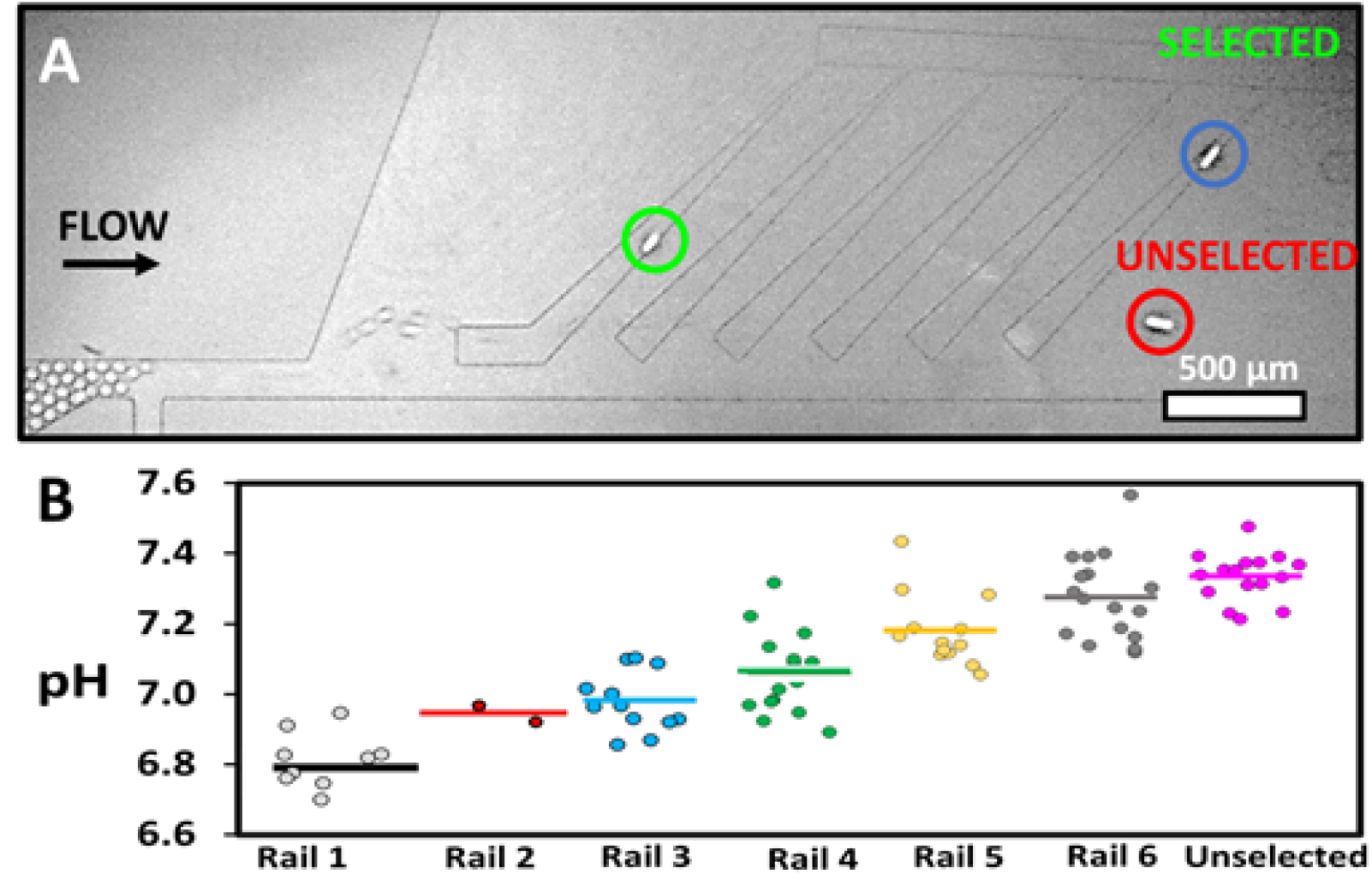
Multirail sorting of droplets. a) A droplet containing a high metabolism cell (circled in green), at pH 6.8, is selected by Rail 1. A droplet containing a lower metabolism cell (circled in blue) at pH 7.2, is selected by Rail 6. Droplets with very low metabolism cell are Unselected (circled in red). Empty droplets are also directed to the Unselected chip exit. b) Successive droplet pH at incubator for droplets selected by each rail and Unselected cell population. Horizontal bar represents average pH value.

A ratiometric pH sensor, pyranine, was used to determine the pH of all droplets after incubation to correlate the pH to droplet trajectory (Figure 3b). The first rail, positioned approximately 1 mm downstream in the sorting region, was found to select droplets with the very lowest pH after incubation, with an average pH of 6.79 ± 0.01 (average ± standard deviation of the mean). These droplets would have the highest interfacial tension when they reach the first rail. The first rail would thus only select cells with the very highest glycolysis levels. Subsequent rails would select droplets stepwise with increasing pH after incubation corresponding to an average pH of 6.94, 6.97, 7.01, 7.17, 7.25 respectively for rails 2 through 6. All these pH measurements have a standard deviation of the mean of approximately 0.01 pH units. Empty droplets, or droplets containing cells with low metabolism, would have the highest pH and hence lowest interfacial tension. These droplets, with an average pH of 7.32 ± 0.01, are not diverted by any rail and flow towards the Unselected outlet. The pH selection of a rail depends on the flow rates in the channel through the Oil Entrainment Inlet (Figure S1),^19^ offering a user-defined parameter for modulating droplet selection. There is overlap of droplet pH between droplets selected by adjacent rails. Part of this overlap can be attributed to the uncertainty of the pH determination, that have a standard deviation of around ± 0.05 pH units based on the variability of pH readings of droplets of identical pH.

This experiment shows that the downstream position of the rail can be used to select cell populations based on the pH change during glycolysis. However, this particular chip geometry does not collect cells into multiple populations as all selected droplets are directed to the same outlet. Rail spacing and shape can be optimized to direct distinct glycolytic cell populations towards separate outlets.

Figure 4a and supplemental video S3 show the sorting of cells into three populations with low, middle and high glycolysis. Here, rail spacing has been doubled to 600 μm to minimize the overlap of droplet pH between adjacent rails. The first rail spans the channel width and utilizes a horizontal rail section to direct droplets to the top outlet of the device. The second rail directs droplets to a middle outlet. Lastly, the Unselected outlet comprises the bottom portion of the sorting region. A rail directed in the downward direction ensures that any droplets that may undergo a slight deviation in path are still directed towards the Unselected outlet. Rail position and design are provided in Figures S2 and S3, respectively.

**Figure 4.**
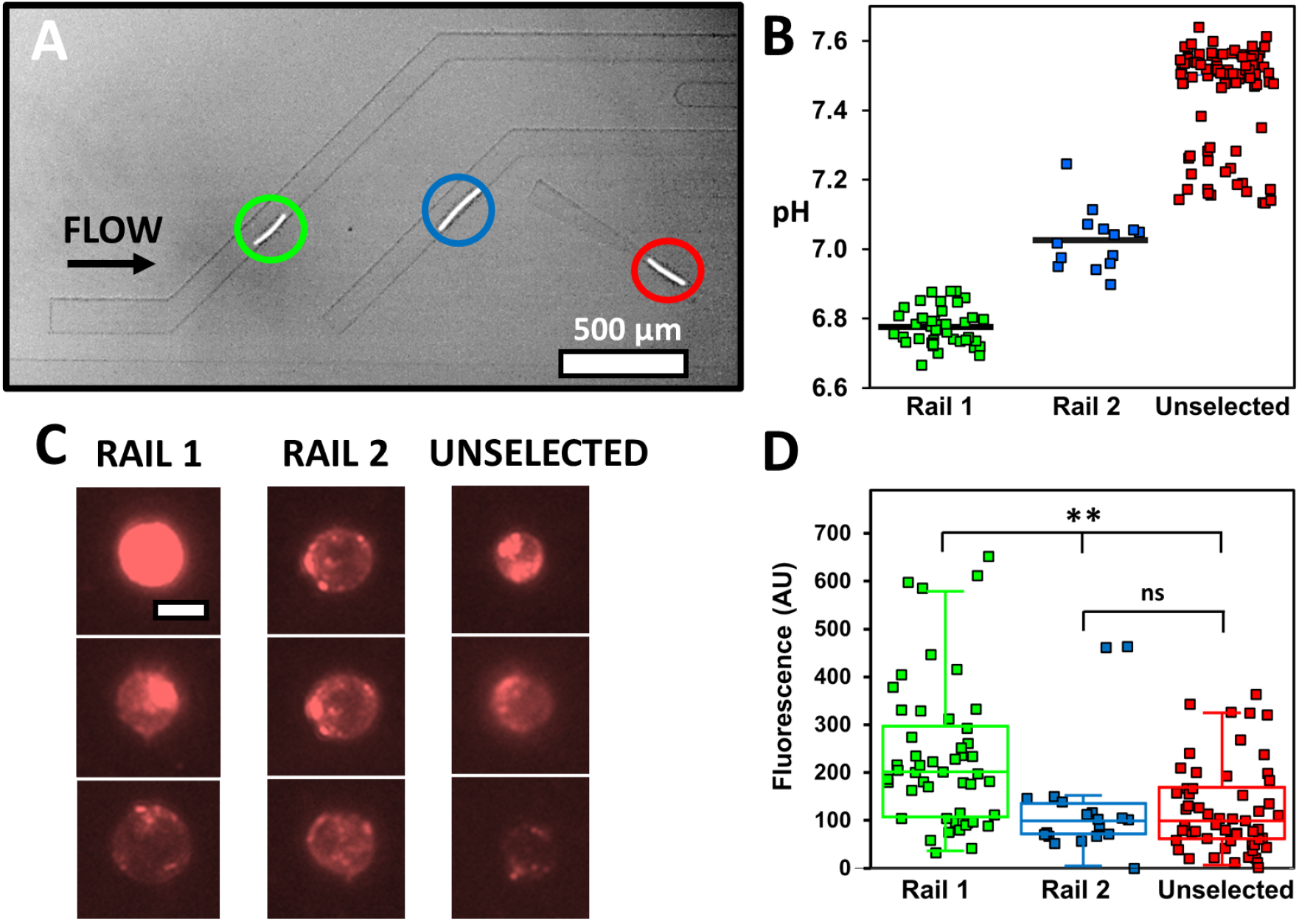
Correlation of glycolysis of T-cells to activation markers a) A droplet containing a high metabolism cell (circled in green), is selected by Rail 1 to top outlet. A droplet containing a lower metabolism cell (circled in blue), is selected by the second rail to middle outlet. Droplets with very low metabolism (circled in red) are directed to the Unselected outlet. Empty droplets are also directed to the Unselected outlet. Streaks are due to fast movement of droplets containing a cell marker. b) pH of droplets selected by each rail and Unselected cell population. Horizontal bar represents average pH value. c) Representative fluorescence images of cells with CD69 activation marker for cells collected from Rail 1, Rail 2 and Unselected. Scale bar is 10 µm. d) Fluorescence intensity of CD69 activation marker. Middle horizontal line represent median while lower and upper boxes represent 25 and 75^th^ percentile. Whiskers shows range of data. **\*\*** indicates p < 0.01 and ns is for non-significant.

This device was used to separate activated T-cells (Jurkat cells) based on three distinct glycolysis levels and, for the first time, correlate each level with surface activation markers (CD69) at the single-cell level. In a previous paper, we showed that SIFT can isolate activated T-cells from naive cells based on their elevated glycolysis.^23^ Upon activation by antigen-presenting cells or by antibody-coated beads that mimic their activation complexes, T-cells rapidly ramp up glycolysis. This metabolic reprogramming supports rapid proliferation and differentiation.^11,35^ Activated T-cells also display distinctive surface markers over the course of hours and days after activation, such as CD25, CD69 and CD71.^36^ The strength of activation has been previously correlated to extracellular acidification rate.^37^ Strength of activation favors different differentiation outcomes^38^ and apoptosis sensitivity.^39^ Glycolysis levels were compared to the presence of the quickly upregulated surface activation marker CD69 to determine if the two are distinct or complimentary indicators of activation. In other words, whether the SIFT device, which sorts based on glycolytic activity, isolates a novel population of cells compared to conventional markers of activation.

Figure 4b shows the droplet pH of the three populations sorted based on glycolytic level (Figure 4b). The first rail sorts droplets with the lowest pH after incubation with an average pH of 6.78 ± 0.01. Droplets selected by the middle population and hence intermediate glycolysis levels have an average pH of 7.03 ± 0.01. Lastly the droplets collected in the Unselected outlet contain droplets of the highest pH, above 7.15. The unselected droplets contain a cluster of droplets near pH 7.5 and another centered around pH 7.2. These experiments were performed without fluorescent cell markers to avoid spectral interference with the fluorescent activation marker. This makes it difficult to confirm the presence of a cell in a fast moving droplet. However, from comparable experiments containing cell markers, the population of droplets at pH 7.5 would be mostly empty droplets while the droplets around 7.2 would contain cells with low glycolysis.

A logistical regression was used to estimate the threshold of selection between different rails (Figure S7). Between rail 1 and rail 2 the threshold was determined to be a pH of 6.89. There is no overlap between the two populations so an error could not be determined directly from the fit. In this case, an uncertainty would be estimated by the closest points between the two populations. This would lead to a selection threshold of pH 6.89 ± 0.01. In the case of rail 2 and Unselected the threshold was determined to be 7.11 ± 0.06 (95% confidence interval). The errors in sorting thresholds confirm that accurate sorting is achievable with three populations with errors that are comparable to those observed in binary SIFT sorting.^19,23^

After sorting and isolation, the CD69 levels were quantified by fluorescence for the three populations of T-cells labelled with anti-CD69. A protocol was developed to ensure recovery and viability of cells after on-chip sorting (described in detail in the Materials and Methods and Figure S4). Representative fluorescence images of the three glycolysis populations are provided in Figure 4c. A distribution of fluorescence intensities is observed for all three populations.

Figure 4d shows the average fluorescence intensity and box plot for individual cells for the three sorted cell populations. Cells collected from rail 1, with the highest glycolysis level, includes cells with the highest fluorescence intensity. This population displays the largest mean and is statistically different when compared to the cell population isolated from rail 2 and Unselected outlets (p < 0.01). Rail 2 and the Unselected cells show lower fluorescence and were not statistically different (p > 0.05). A duplicate dataset came to the same conclusions (data not shown).

The strength of activation is expected to promote higher glycolysis and activation maker expression. Thus, a correlation between glycolysis level and the activation markers may be expected. Rail 1 isolated the cells with the most intense CD69 signal. However, the three populations showed overlapping ranges of CD69 intensities, with the middle glycolysis population undistinguishable from the low glycolysis population. This suggests a more complex relationship between the marker intensity and glycolysis.

## Conclusions

We present a facile and robust method that uses multiple rails for the stepwise isolation of cells based on glycolytic activity. The technique leverages dynamic droplet acidification from surfactant adsorption that leads to a concurrent increase in interfacial tension. It is a passive technique that uses no labels or active components, reducing both cost and complexity. It is particularly well suited to separate cells of the same type that may not have other distinguishing features. Dead cells can complicate or distort downstream analysis, such as RNA-seq.^40^ The device can also be used to exclude these non-viable cells while separating low and high glycolysis cells. The chip presented was designed for the separation of three populations of cells based on glycolysis. However, the number of populations sorted can be expanded by the placement of additional rails and chip outlets. Sorting can be further tuned by controlling the flow speeds, buffer concentration and surfactant concentration. The workflow presented here allows for the recovery and analysis of live sorted cells. The glycolysis level was correlated to CD69 activation markers, demonstrating a proof-of-concept correlation made possible by the technology.

Interestingly, although rapid CD69 surface expression in activated T-cells requires de novo RNA and protein synthesis, cytoplasmic pools of CD69 also exist in resting cells which are independent of this regulation.^41,42^ Thus, the correlation in glycolysis and CD69 expression may not be linear as observed in our results. Glycolysis, however, is now appreciated not only as a byproduct of activation but rather a key regulator of both T-cell activation and differentiation.^14^ Therefore, these results suggest an opportunity to isolate cell populations by a criteria that is largely independent of activation markers. Finally, as glycolysis is also linked to disease state (ie: cancer cells) or activation state (T-cells), the technology enables label-free isolation for study and use of important cell subpopulations for applications in oncology, immunology and immunotherapy.

## Acknowledgments

We acknowledge Brody Sandel of the Biology Department at Santa Clara University for the logistical regression analysis. We would also like to thank Ian Carter-O’Connell from Santa Clara University and Angga Perima from the Pasteur Institute for helpful discussions. TM acknowledges generous funding from the Beckman Scholar program. AG acknowledges support from the Inclusive Excellence Postdoctoral Program at Santa Clara University. AT and CC acknowledge funding from the Clare Boothe Luce Scholar program and KA acknowledges funding from the Bastiani award. KV was supported by the Breakthrough T1D (formerly the Juvenile Diabetes Research Foundation) Advanced Postdoctoral Fellowship. PA is supported for this project by a National Science Foundation Career Award, Grant Number 1751861, the National Institutes of Health grant 1R15GM129674-01 and the Henry Dreyfus Teacher-Scholar Awards Program.

