## Supplemental Info for "Stepwise Isolation of Diverse Metabolic Cell Populations Using Sorting by Interfacial Tension (SIFT)"

### Supplemental Information

**Table of Contents:**

**Video Caption………………………………………………………...………..…...Page S-2**

**Figure S-1…...………………………………………………………………..…..Page S-3**

**Figure S-2…...………………………………………………………………..…..Page S-4**

**Figure S-3…...………………………………………………………………..…..Page S-4**

**Figure S-4…...………………………………………………………………..…..Page S-5**

**Figure S-5…...………………………………………………………………..…..Page S-6**

**Figure S-6…...………………………………………………………………..…..Page S-7**

**Figure S-7…...………………………………………………………………..…..Page S-8**

**Table S-1…...……………………………………………………………..…......Page S-9**

**Video Caption:**

**Video S1. SIFT Sorting with pH Color Indicator:** This color video illustrates the basic principle of SIFT. A pH indicator enables the estimation of droplet pH based on color (color scale provided on top). The video pans in the direction of droplet flow. Cells are encapsulated in droplets and flow through a long serpentine channel. Some droplets contain cells that secrete protons via glycolysis, causing a pH change (evident by a color shift to orange in some droplets). The droplets with low pH (high surface tension) follow a diagonal rail at the end of the chip, while those with high pH (low surface tension) containing no cells or cells with low glycolysis flow horizontally.

**Video S2. Multi-Rail Sorting:**
Multiple rails, positioned at different downstream locations, deflect droplets with varying pH and, consequently, different cell glycolysis levels.

**Video S3. Three Population Cell Sorting:**
In this video, T-cells are directed by rails to three different chip outlets, enabling the collection of cells with varying glycolysis levels: low, medium, and high.

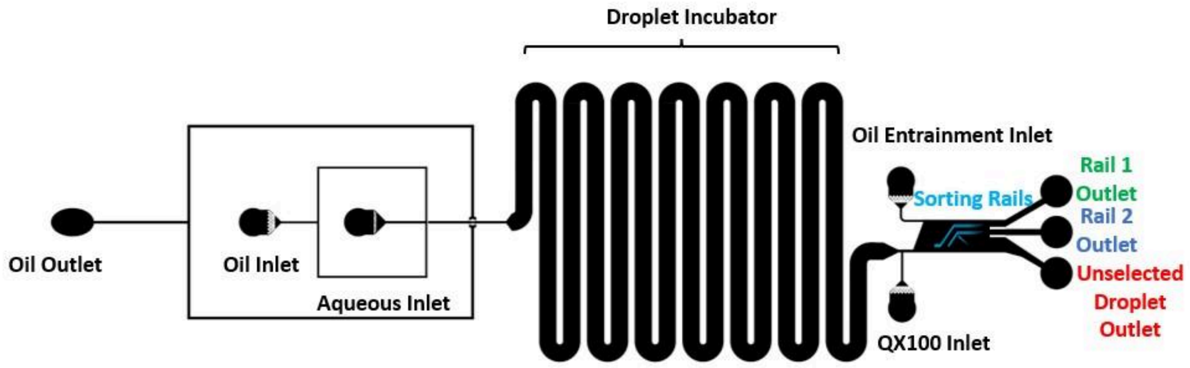


**Supplemental Figure S1. SIFT device channel geometry.**

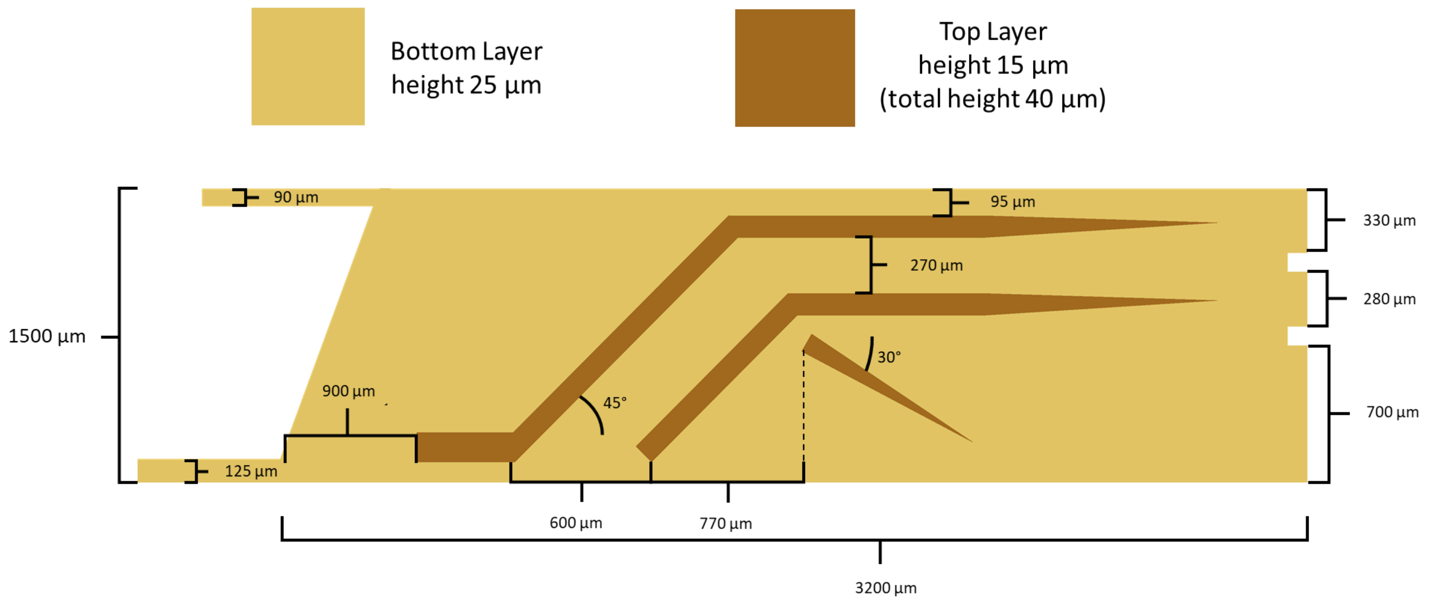


**Supplemental Figure S2. Sorting rail position.** Exact position of rail is approximate as layers are positioned by eye.

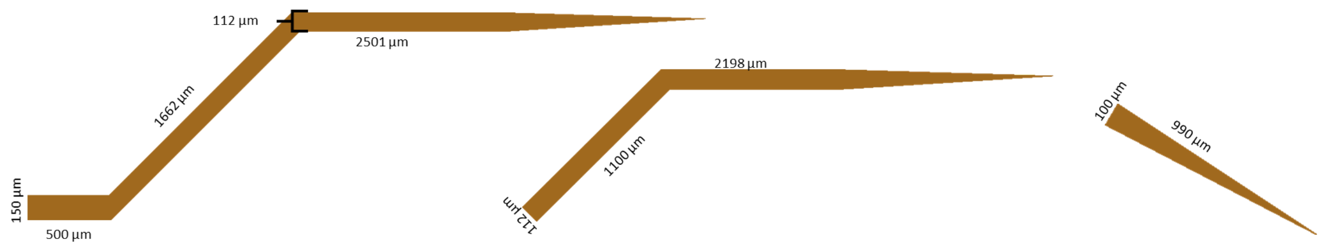


**Supplemental Figure S3. Sorting rail dimensions.**

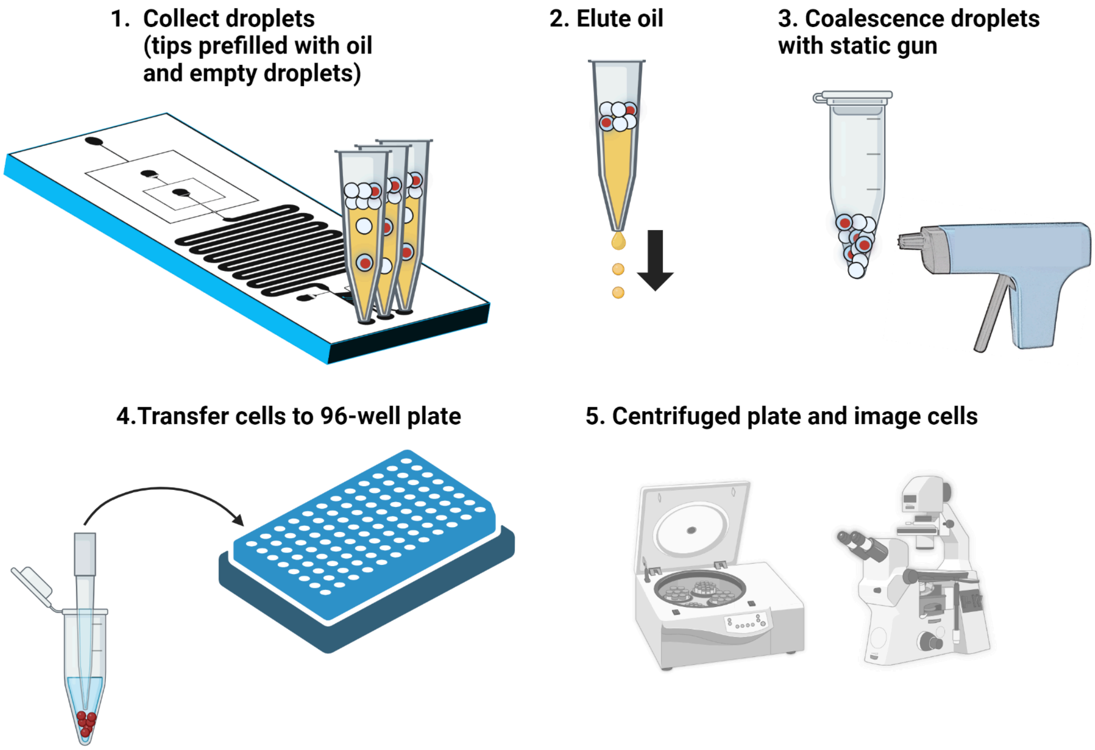


**Supplemental Figure S4.** Workflow for cell collection. Created in BioRender.com.

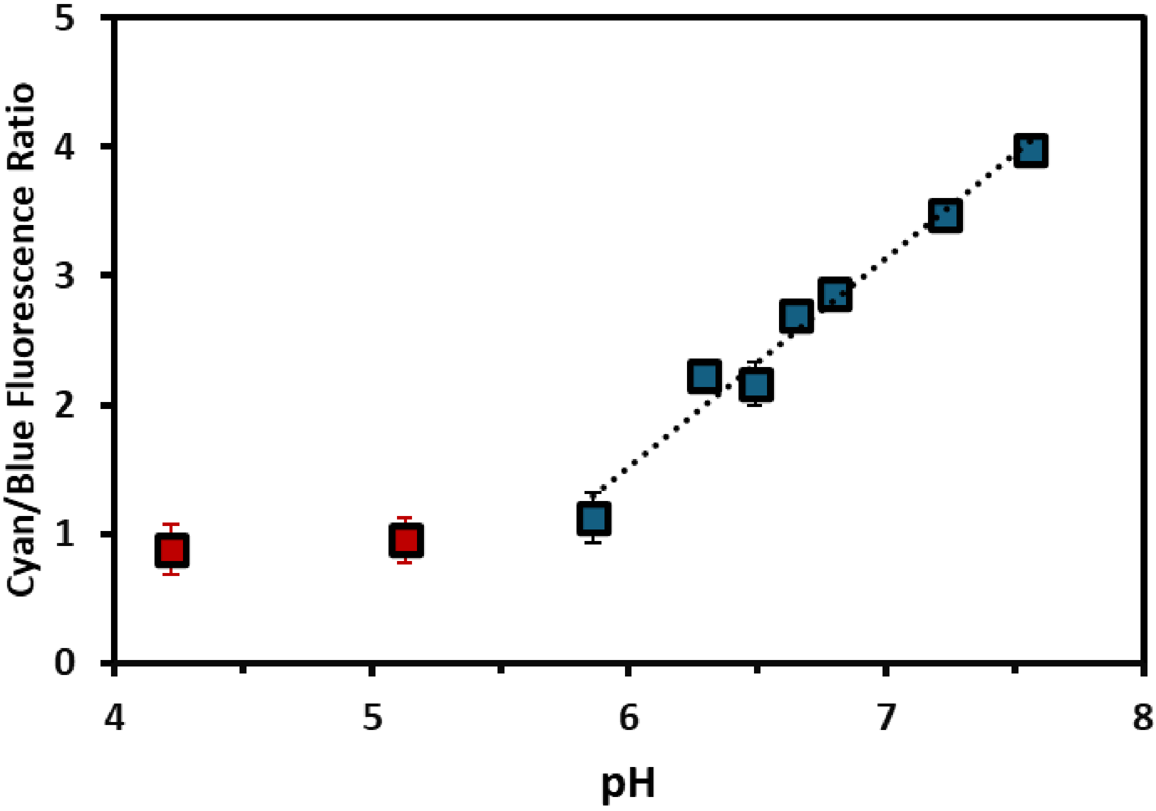


**Supplemental Figure S5.** Cyan to blue fluorescence intensity ratio vs pH of droplets containing fluorescein, a pH sensitive ratiometric fluorescent probe. Solutions of known pH were injected into the microfluidic device and droplets were analyzed for their normalized fluorescence intensity through excitation with cyan (479 nm) and blue (440 nm) light. The calibration curve was obtained under the same conditions as cellular experiments. The slope of the calibration curve was determined from the linear fit for points from pH 5.8 to 7.6. The y-intercept was adjusted for each experiment using the known pH of empty droplets. Error bars represent the standard deviation of the average cyan/blue ratio of five representative droplets for each pH measured and are not visible when smaller than the marker.

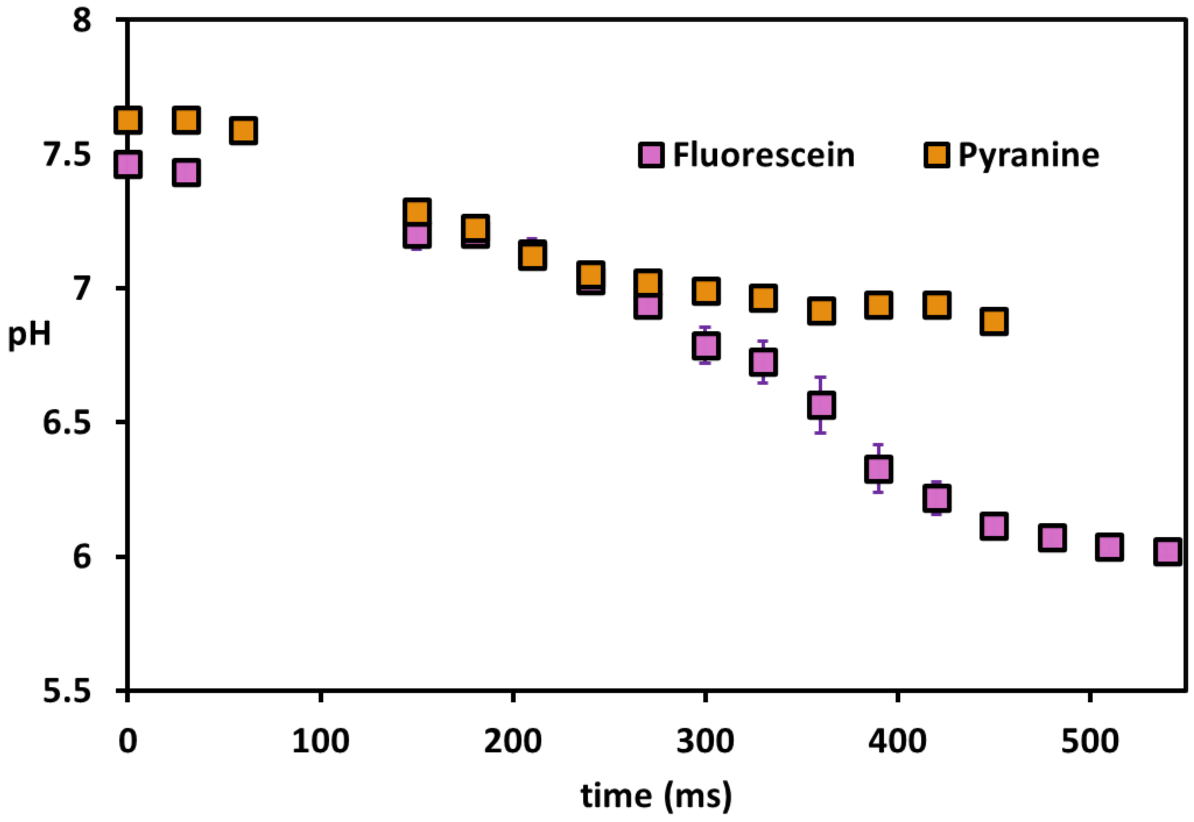


**Supplemental Figure S6.** Comparison of acidification over time of the droplets through the adsorption of surfactant measured by pyranine and fluorescein. The initial pH of droplets measured by fluorescein or pyranine are similar and follow similar dynamics till 300 milliseconds. The measurements based on pyranine however do not decrease below around pH 7.0, near the minimum pH measurable with this probe. In contrast, measurements based on fluorescein show a droplet pH that decreases to around pH 6.0. Error bars represent the standard deviation of the mean and are not visible when smaller than the marker.

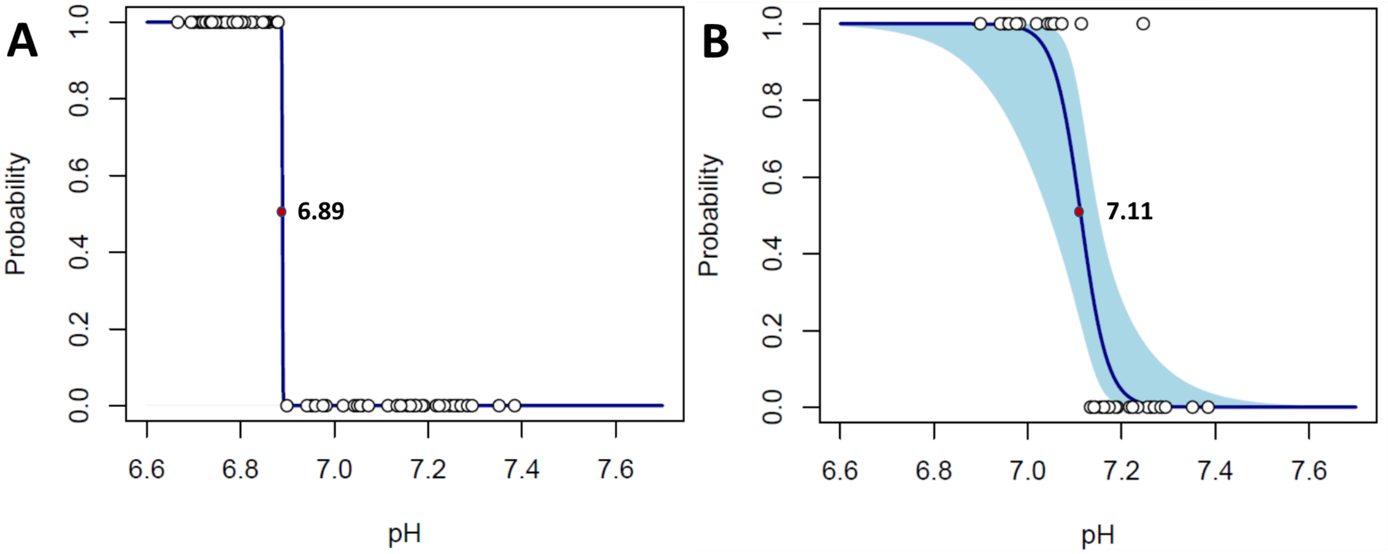


**Supplemental Figure S7. Logistic Regression Fits** (**A**) Logistic regression fit of selection by Rail 1 vs. Rail 2 or Unselected (**B**) Logistic regression fit for Rail 2 vs. Unselected. pH thresholds are indicated on graph and represent where there is equal probability that droplets are selected or unselected. The 95% confidence limit is indicated in light blue.

**Supplemental Table S1. Typical flow parameters**. Channel geometry is provided below for reference. Negative flows below are opposite in direction to the main flow in the channel.

| Inlets and Outlets | Flow Rates (μL/min) |
|---|---|
| Aqueous Inlet | 0.3 - 0.5, most commonly 0.3 |
| Oil Inlet | 3 – 5, most commonly 3 |
| QX100 Inlet | 5 – 10 |
| Oil Entrainment Inlet | 25 – 40 |
| Oil Outlet | - 2 to -4 |

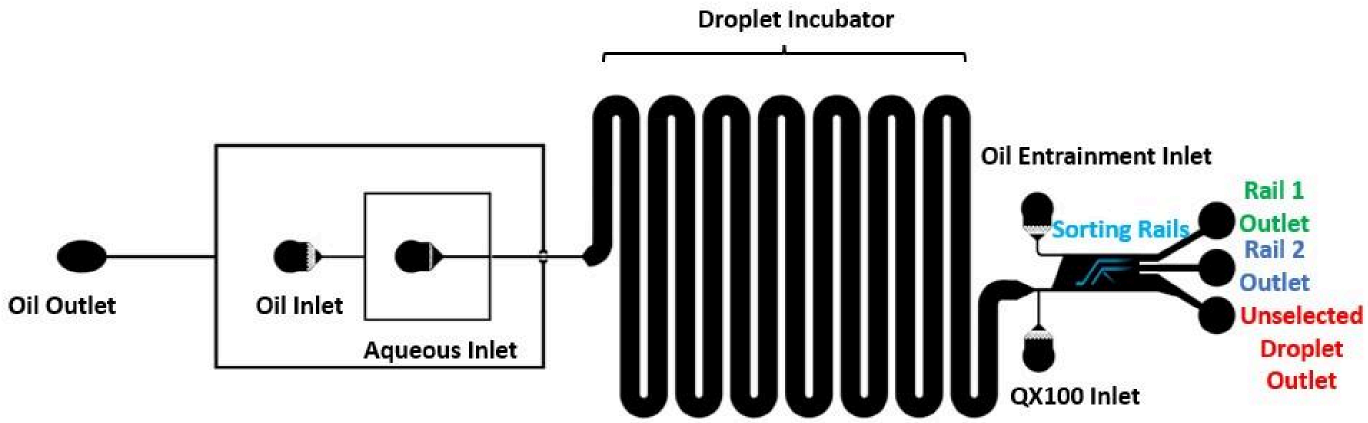
